# Immediate myeloid depot for SARS-CoV-2 in the human lung

**DOI:** 10.1101/2022.04.28.489942

**Authors:** Mélia Magnen, Ran You, Arjun A. Rao, Ryan T. Davis, Lauren Rodriguez, Camille R. Simoneau, Lisiena Hysenaj, Kenneth H. Hu, The UCSF COMET Consortium, Christina Love, Prescott G. Woodruff, David J. Erle, Carolyn M. Hendrickson, Carolyn S. Calfee, Michael A. Matthay, Jeroen P. Roose, Anita Sil, Melanie Ott, Charles R. Langelier, Matthew F. Krummel, Mark R. Looney

## Abstract

In the severe acute respiratory syndrome coronavirus 2 (SARS-CoV-2) pandemic, considerable focus has been placed on a model of viral entry into host epithelial populations, with a separate focus upon the responding immune system dysfunction that exacerbates or causes disease. We developed a precision-cut lung slice model to investigate very early host-viral pathogenesis and found that SARS-CoV-2 had a rapid and specific tropism for myeloid populations in the human lung. Infection of alveolar macrophages was partially dependent upon their expression of ACE2, and the infections were productive for amplifying virus, both findings which were in contrast with their neutralization of another pandemic virus, Influenza A virus (IAV). Compared to IAV, SARS-CoV-2 was extremely poor at inducing interferon-stimulated genes in infected myeloid cells, providing a window of opportunity for modest titers to amplify within these cells. Endotracheal aspirate samples from humans with the acute respiratory distress syndrome (ARDS) from COVID-19 confirmed the lung slice findings, revealing a persistent myeloid depot. In the early phase of SARS-CoV-2 infection, myeloid cells may provide a safe harbor for the virus with minimal immune stimulatory cues being generated, resulting in effective viral colonization and quenching of the immune system.

---

The SARS-CoV-2 pandemic has led to over 6 million deaths worldwide. Respiratory viruses like SARS-CoV-2 are known to infect and replicate in airway epithelial cells, inducing lung injury with often fatal outcomes^1^. The access to large numbers of human samples has given the opportunity to extensively study immune responses to COVID-19^2–4^. Unfortunately, once patients are hospitalized, the host-pathogen responses have been in progress for days or weeks, creating a challenging problem for understanding the very early host responses to SARS-CoV-2 infection in humans. While animal models are useful in this regard, they do not fully recapitulate human complexity, including appropriate expression of relevant ligands. Here, we used precision-cut lung slices (PCLS) obtained from human lungs to study early hostpathogen responses in a system replete with the full repertoire of lung stromal and immune cells.

We have previously developed a model of PCLS in the mouse lung^5,6^ that we have now applied to human lungs donated for research (Supplementary Table 1). A lung lobe was inflated using 2% low melting point agarose and 300 μm PCLSs were produced for tissue culture and direct infection with SARS-CoV-2 (USA-WA1/2020; MOI 0.1 – 1) for up to 72 hours (Fig. 1a). Imaging of the infected PCLS (Fig. 1b-c; non-infected in Extended Data Fig. 1) revealed spike staining in epithelial cells (EpCAM^+^) that were also ACE2 positive (Fig. 1b). PCLSs were enzymatically digested for multicolor flow cytometry analysis (Fig. 1a). To further investigate infection, cells were stained for dsRNA as well as spike (Fig. 1d-e). Epithelial cells (CD45^-^ EpCAM^+^ CD31^-^) displayed both dsRNA and spike signal after 72h of SARS-CoV-2 infection (Fig. 1d and Extended Data Fig. 2a). At 72h post-infection, 4 to 10% of epithelial cells were spike^+^ and 2 to 6% dsRNA^+^. Both imaging and flow cytometry showed that SARS-CoV-2 was able to induce epithelial infection in a small-scale human lung model.

**Figure 1.**
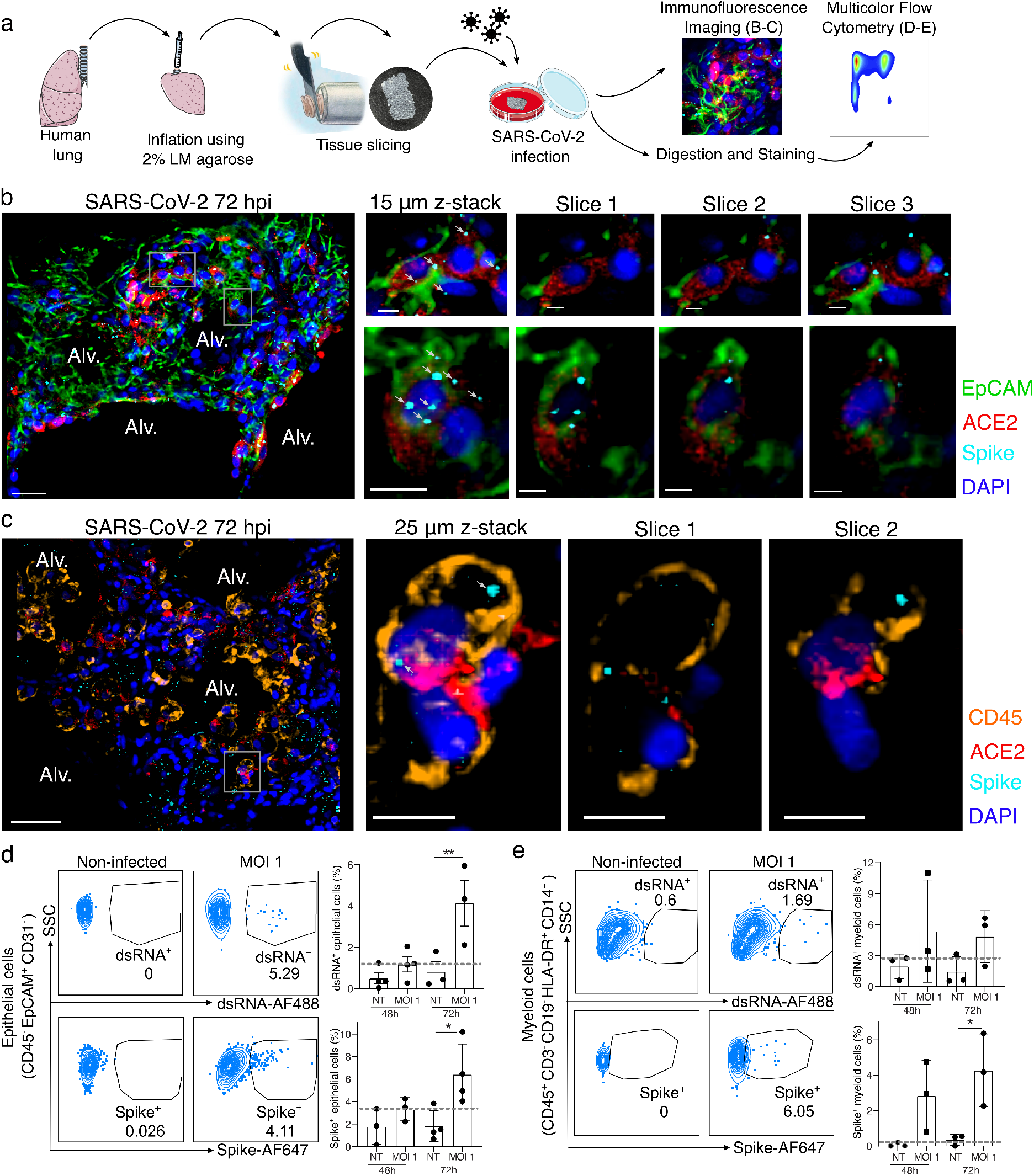
SARS-CoV-2 infects both epithelial and immune cells in human PCLSs. **(a)** Schematic diagram of the experimental design. Human lung lobes were inflated with 2% low-melting point agarose and sectioned into 300 μm precision cut lung slices (PCLS), which were cultured in 24-well plates and infected with SARS-CoV-2 for 48 or 72h. After incubation, PCLS were either fixed and stained for confocal imaging **(b, c)** or dissociated and stained for flow cytometry **(d, e). (b, c)** PCLS were infected with SARS-CoV-2 for 72h at MOI 0.1 or 1 and used for confocal imaging. Alveolar spaces (Alv.) are indicated in the large image (scale bar = 50 μm). Zoom area is marked by the white rectangle. For each zoomed area, 15 or 25 μm z-stacks appears on the side with single x-y sections (scale bar =10 μm). **(b)** PCLS were stained for DAPI (dark blue), EpCAM (green), ACE2 (red) and spike (light blue), (c) PCLS were stained for DAPI (dark blue), CD45 (orange), ACE2 (red) and spike (light blue), **(d, e)** PCLS were infected at MOI 1 for 48 and 72h. PCLS were dissociated and cell suspension was stained for flow cytometry analysis (n=3-4). Infection was assessed by intracellular spike and dsRNA staining in **(d)** epithelial and **(e)** myeloid cells. Grey dashed lines indicate detection limit of assays. Data are mean ± SEM. Each dot represents the average percentage of dsRNA÷ or spike÷ myeloid cells of 2 to 3 individual lung slices from one donor, *p<0.05, **p<0.01.

Using both imaging and flow cytometry, we also observed spike and dsRNA signal in lung immune cells (Fig. 1c and e), the former prevalent from the earliest 48-hour timepoint. Spike was colocalized to CD45^+^ ACE2^+^ cells (Fig. 1c; non-infected in Extended Data Fig. 1b) and similarly dsRNA was found in these cells (Extended Data Fig. 2b), supporting that concept that immune cells may either be infected by SARS-CoV-2 or phagocytose the virus. Flow cytometry allowed us to further characterize spike^+^ and dsRNA^+^ immune cells (Fig. 1e). We observed significant dsRNA and spike signal in lung myeloid cells (CD45^+^ CD3^-^ CD19^-^ HLA-DR^+^ CD14^+^cells, including interstitial macrophages, monocytes and monocyte-derived dendritic cells^7^) at 48-72h post-infection. These data support that immune cells, particularly myeloid cells, have profound and early interactions with SARS-CoV-2, potentially leading to a major role in shaping the immune response.

To further characterize the transcriptional influence of SARS-CoV-2 infection upon specific cell populations, during the first days of exposure, we applied single-cell RNA sequencing (scRNA-seq) analysis to cells obtained at various timepoints after PCLS infection (Fig. 2). Hierarchical analysis identified clusters representing highly heterogenous lung complexity, including eight populations of non-immune cells, lymphocytes (T cells, B cells, NK), and four populations of myeloid cells (Fig. 2a and Extended Data Fig. 3a)^8^. To appreciate unique changes in lung composition and gene expression induced by SARS-CoV-2, we compared lung slices infected by either SARS-CoV-2 or Influenza A virus (IAV) (Fig. 2b). In response to IAV, the most profound change was a reduction in lung fibroblast and epithelial cell proportions, consistent with previous reports^9^. SARS-CoV-2, in contrast, produced no significant trends in lung non-immune cell populations, compared to controls. IAV was similarly destructive in the immune populations, producing a notable overall decrease in cells clustered in “Myeloid 1” (Extended Data Fig. 3a). Conversely, SARS-CoV-2 infection actually increased the myeloid fraction over time, increasing by almost 50% relative to controls at 72h (Fig. 2b).

**Figure 2.**
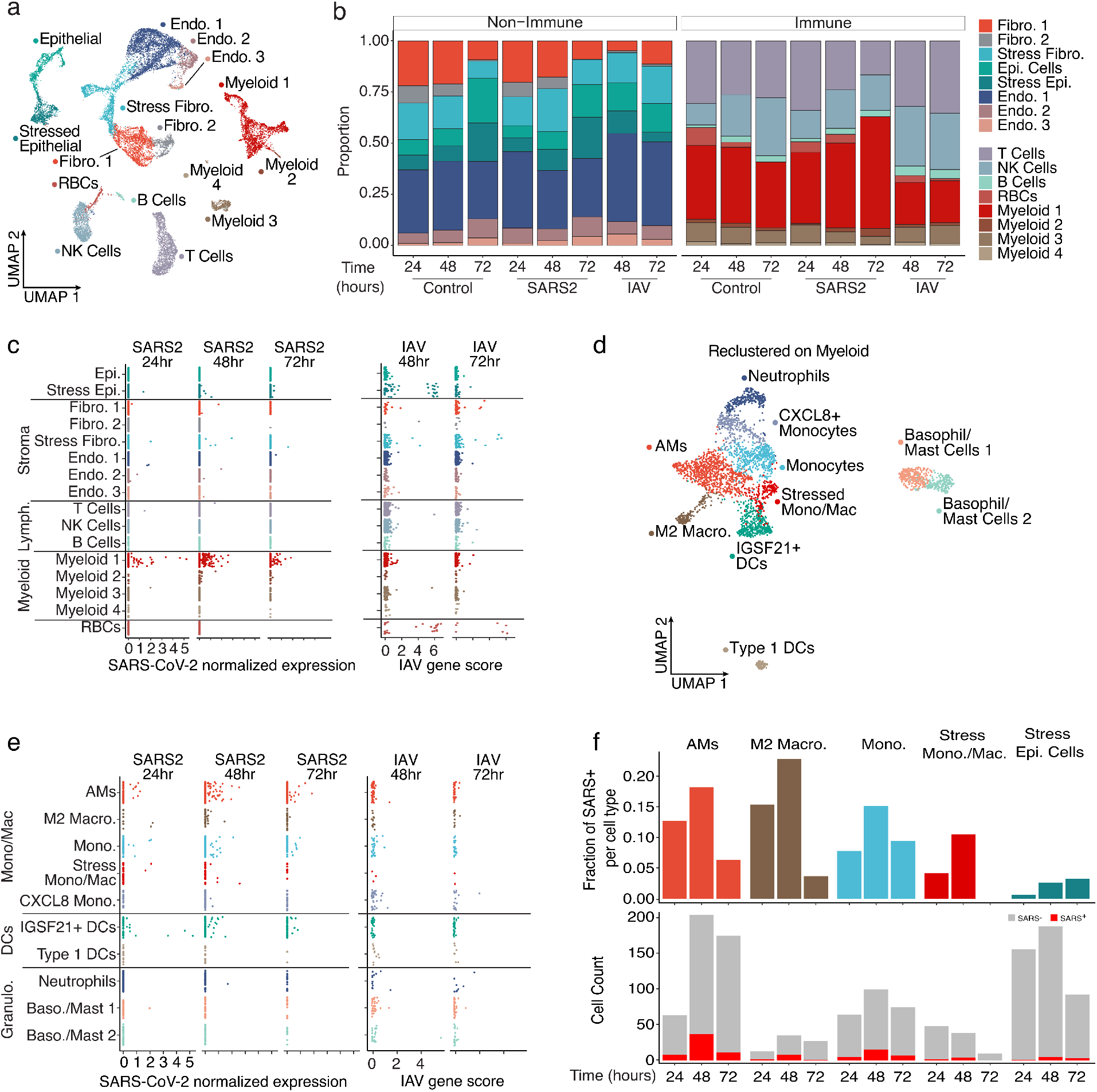
SARS-CoV-2 displays tropism for myeloid cells compared to IAV. **(a)** A Uniform Manifold Approximation and Projection (UMAP) visualization of cells from control, SARS-CoV-2, and lAV-infected PCLS collected at distinct times, (b) Relative quantification of cell types from the different experimental conditions, stratified by timepoint, **(c)** Scatterplots describing the library-normalized SARS-CoV-2 expression across various cell types in SARS-CoV-2-infected PCLS (left) or IAV gene score in lAV-infected PCLS (right), **(d)** A UMAP of finely annotated myeloid cell types in the dataset, **(e)** Distribution of infected myeloid cells similar to **(c). (f)** The fraction (top) and numbers of SARS-CoV-2-positive cells (bottom) at different timepoints.

Aligning the scRNA-seq data on the two viral genomes revealed that the main targets for IAV infection were epithelial cells and fibroblasts (Fig. 2c), consistent with the observed loss of these populations in PCLSs (Fig. 2b). In addition, IAV reads were also sporadically distributed in several other populations including immune cells. This may be attributable to the IAV entry receptor (sialic acid) being widely distributed on the surfaces of many cell types or perhaps to phagocytosis. SARS-CoV-2 reads by comparison were distinctly enriched in myeloid cells, even at the earliest 24h timepoint. Only a few reads were identified in non-immune cells (Fig. 2c). To identify the subpopulation of myeloid cells targeted by SARS-CoV-2, the four myeloid clusters from Fig. 2a were combined and re-clustered, resulting in the definition of ten myeloid populations that included neutrophils, dendritic cells (DCs) and multiple subpopulations of monocytes and macrophages (Fig. 2d and Extended Data Fig. 3b). IAV reads remained sporadic across all 10 clusters (Fig. 2e). Conversely, SARS-CoV-2 tropism among these clusters was more selective with alveolar macrophages (AMs), IGSF21^+^ DCs, and monocytes accounting for the majority of significant viral reads (Fig. 2e). To account for the variable frequency of each population in Fig. 2e, we analyzed the SARS-CoV-2 read frequency and plotted this over time. This demonstrated that viral reads rose in unison in both myeloid and epithelial cell populations from 24 to 48h and then decreased in the myeloid populations at 72h (Fig. 2f). This could be the result of resolution of the infection in the PCLS, or an exhaustion of host cells targeted by the virus.

To investigate how SARS-CoV-2 affects human lung myeloid cells, we focused on the dominant AMs, which are also anatomically within the airspaces and so directly exposed to virus^10^. A bronchoalveolar lavage (BAL) in the donor lungs (Fig. 3a) produced a sample enriched in AMs, defined here as CD45^+^ CD169^+^ HLA-DR^+^ (Extended Data Fig. 4a, d-e)^11^ and comprising notably few epithelial cells (<1% of live cells) (Extended Data Fig. 4b-c). AMs were analyzed for ACE2 expression (Fig. 3b), and we observed variability amongst lung donors with on average ~10-20% of ACE2^+^ AMs, but with one donor at ~80% (Fig. 3c). We incubated the cell population with SARS-CoV-2 and then analyzed cells by flow cytometry (Fig. 3a, d-e; Extended Data Fig. 4a, f-g). After 48h with SARS-CoV-2 at MOI 0.1, we detected spike in 1-10% of the AMs (Fig. 3d, Extended Data Fig. 4a). Somewhat surprisingly, an MOI of 1 did not significantly increase spike^+^ AM percentage compared to MOI 0.1 (Fig. 3d), suggesting that either cells were somehow protected at higher titers—perhaps due to increase antiviral sensing and subsequent ISGs—or that a plateau was reached in the number of cells that were capable of being infected. At 48h, viability of AMs was high in both MOI groups, an effect that suggests the virus was not inducing AM cell death (Extended Data Fig. 4f) contrary to observations in blood monocytes from COVID-19 subjects^12^. However, of the spike^+^ AMs, the majority were ACE2^+^ (Extended Data Fig. 4g) pointing to a specific but not obligate role for ACE2 in licensing AM viral entry. In support of this, when we used an ACE2 blocking antibody^13^ incubated with AMs 2h prior to SARS-CoV-2, we always observed a significant decrease in spike^+^ AMs (Fig. 3e) but this was rarely complete, even despite using saturating concentrations of the ACE2 antibody. Taken together, these data indicate that SARS-CoV-2 entry into AMs was significantly mediated by ACE2 expression and did not require epithelial cells as an intermediate host, since our BAL preparation lacked these cells.

**Figure 3.**
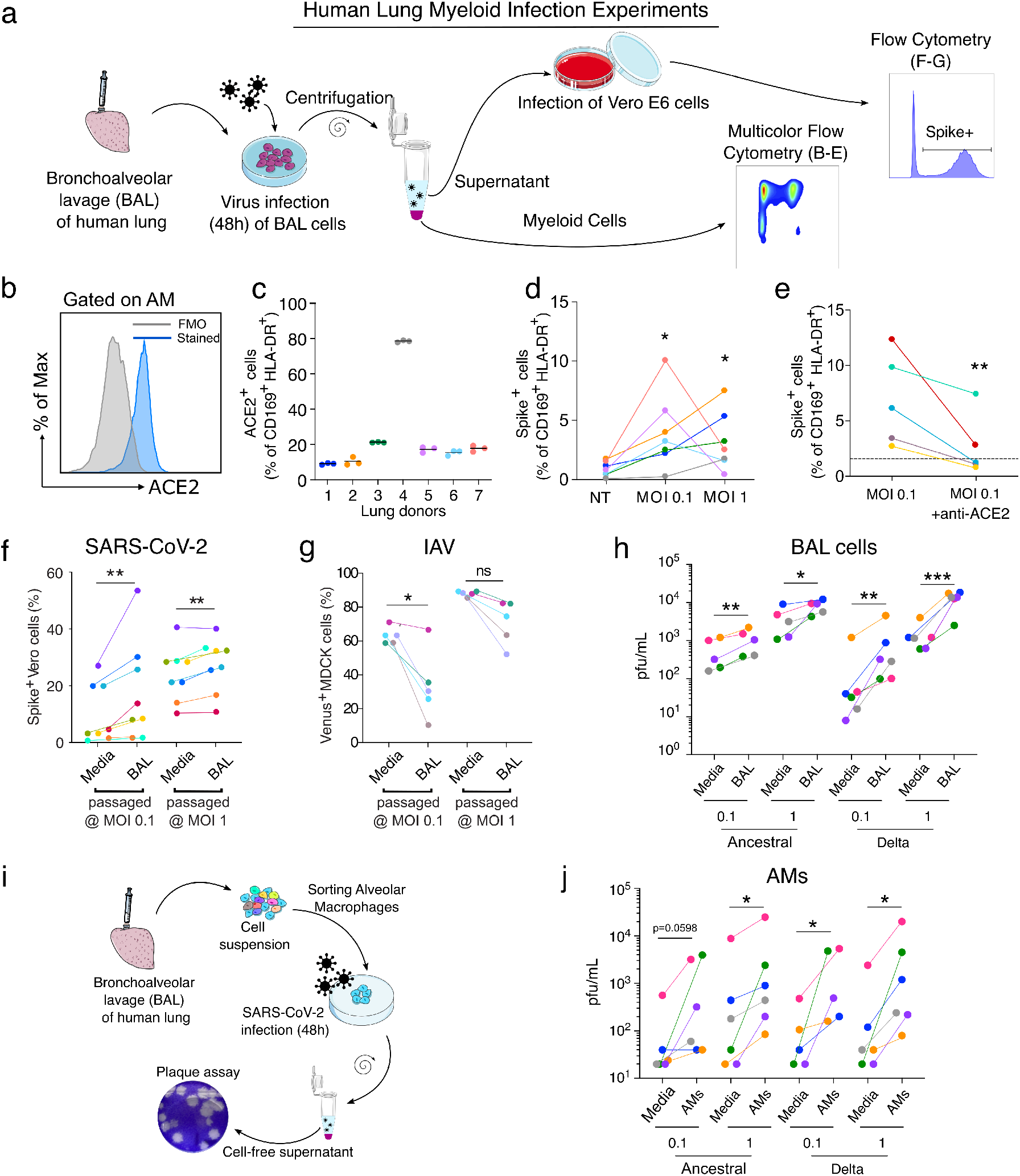
SARS-CoV-2 infection of alveolar macrophages is ACE2-dependent and amplifies viral titer, **(a)** Bronchoalveolar lavage (BAL) of human lungs yields cells that were infected with SARS-CoV-2 for 48h. Infected cells were analyzed by multicolor flow cytometry. Supernatant of infected BAL cells was used to infect Vero E6 cells. 24h after infection, infected cells were quantified by flow cytometry, (b) ACE2 protein expression was assessed on alveolar macrophages (AMs: CD169+ HLADR+ cells) by flow cytometry, (c) Proportion of ACE2+ AMs was measured (n=7). **(d)** AM infection was measured by flow cytometry using spike staining (each color represents a human lung donor, n=7). **(e)** ACE2 blocking antibody was added to BAL cells before infection (MOI 0.1). Cells were analyzed by flow cytometry (n=5). (f) BAL cells were infected with SARS-CoV-2 (BAL passaged). As a control, SARS-CoV-2 was incubated in culture media alone (Media). Cell-free supernatant was used to infect Vero E6 cells. At 24h after infection, Vero E6 cells were stained for intracellular spike expression and analyzed using flow cytometry (n=8). (g) Similarly, lAV-Venus was used to infect BAL cells or incubated with media. Cell-free supernatant was used to infect MDCK cells. At 24h after infection, Venus expressing MDCKs percentage was measured by flow cytometry (n=5). **(h)** Plaque assay was used to further assess viral titer in supernatant of infected BAL cells (n=5-6). (i) AMs were sorted from BAL samples, (j) Following 48h of SARS-CoV-2 infection (Ancestral, delta), viral titer was determined by plaque assay (n=5-6). ns=not significant, *p<0.05, **p<0.01, ***p<0.001

Macrophage infection by viruses such as IAV has long been described as abortive, but several studies have shown that the IAV H5N1 strain is capable of virus production in macrophages^14^. To study the ability of AMs that were exposed to virus to produce and release new viruses, SARS-CoV-2 and IAV (fluorescent strain, PR8-venus) were incubated with BAL cells (MOI 0.1 or 1) as above and the resulting titer in the media was then re-assessed. As a control, the same quantities of viruses were incubated only with media. After 48h of incubation, cell-free supernatant was then collected from the control (media) and BAL groups and used to infect Vero E6 (SARS-CoV-2) or MDCK cells (IAV). After 24h, the infection of these readout cells was assessed by spike staining and flow cytometry (Fig. 3f-g, Extended Data Fig. 5a-b). By quantifying Vero E6 cells that are positive for spike (SARS-CoV-2) or MDCK cells positive for Venus (IAV), we could quantify the infective load of virus present in cell-free supernatant after incubation with either BAL cells or cell-free media (Fig. 3a). For SARS-CoV-2, incubating the virus with BAL cells amplified virus at both MOI 0.1 and 1 compared to incubating with media alone (Fig. 3f, fold-change in Extended Data Fig. 5a). We observed the opposite effect with IAV, which decreased the amount of virus present at MO1 0.1 (Fig. 3g, fold-change in Extended Data Fig. 5b).

SARS-CoV-2 has been continuously evolving, resulting in the emergence of several variants of concern (VOC), the most pathogenic of which has been the delta variant, which we directly compared to ancestral USA-WA1/2020 in our system. As previously done, BAL cells were infected with SARS-CoV-2 viruses for 48h. To determine SARS-CoV-2 viral production by BAL cells, we used a plaque assay on BAL supernatant (Fig. 3h). BAL cells increased viral titer for both ancestral and delta (Fig. 3h, plaque assay in Extended Data Fig. 5c, fold-change in Extended Data Fig. 5d) confirming productive infection of BAL cells by these variants.

We next sorted AMs from BAL (gated as CD3^-^ CD19^-^ EpCAM^-^ CD45^+^ HLA-DR^+^ CD169^+^) before infection. After 48h infection with SARS-CoV-2 variants (ancestral and delta) at MOI 0.1 or 1, cell-free supernatant was harvested and used for the plaque assay (Fig 3i). Consistent with results from BAL incubations, AM incubation with SARS-CoV-2 induced an increase of viral titer at both MOI 0.1 and 1 (Fig. 3j, plaque assay in Extended Data Fig. 5e, fold-change in Extended Data Fig. 5f). Anti-viral responses (ISG induction) were increased in both variants with increasing MOI most significantly with ancestral (Extended Data Fig. 5g). Taken together, our results show that incubation of AMs with SARS-CoV-2, but not IAV, leads to a productive infection of these cells and viral propagation without cell death.

To compare our findings in the isolated human lung model to clinical COVID-19 cases, endotracheal aspirates (ETA) were sampled from seven intubated subjects with the acute respiratory distress syndrome (ARDS) from COVID-19 (Supplementary Table 2) and analyzed by scRNA-seq (Fig. 4a). Clustering showed that the cellular population was predominantly composed of myeloid cells (Fig. 4a, far right), which contained macrophages, neutrophils, and some DCs (Fig. 4c). Analysis of SARS-CoV-2 normalized expression revealed that SARS-CoV-2 reads localized mainly to macrophages, but in some cases were also found in neutrophils and T cells (Fig. 4b), similar to prior results^15^. The ETA samples were obtained at different times after intubation, but the timing did not correlate with the quantity or SARS-CoV-2 reads (Extended Data Fig. 6). In fact, one subject was sampled at 40 days after intubation but still had detectable viral reads in macrophages, which may point to a long-lasting depot effect. As in the PCLS model, several macrophage subpopulations (Extended Data Fig. 7) were found to have SARS-CoV-2 reads with almost 25% of AMs being virus positive (Fig. 4d).

**Figure 4.**
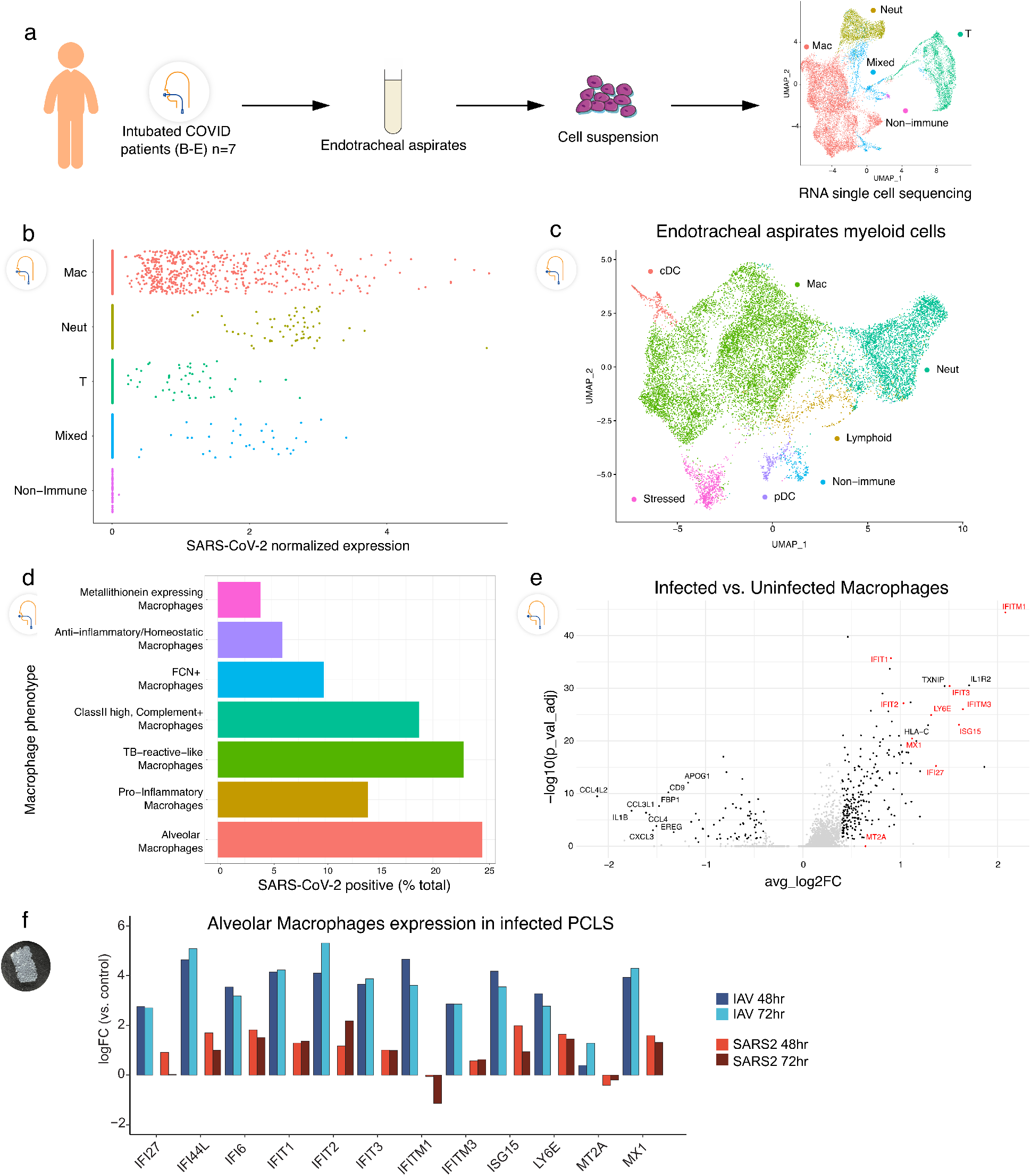
Tropism of SARS-CoV-2 for myeloid cells in endotracheal aspirates from COVID-19 subjects with ARDS, **(a)** Endotracheal aspirates were collected from SARS-CoV-2 infected subjects with ARDS and subjected to scRNA-seq. UMAP at far right shows landmark populations, **(b)** Distribution of per-cell normalized SARS-CoV-2 expression in landmark cell types, **(c)** UMAP projection of myeloid subtypes in endotracheal aspirates, **(d)** Fraction of SARS-CoV-2 positive cells per myeloid cell type **(e)** A volcano plot of SARS-Cov-2 positive AMs vs uninfected AMs. Interferon stimulated genes (ISGs) are highlighted in red. **(f)** Log2 Fold Change of select ISGs in the PCLS experiment in IAV- and SARS-CoV-2-infected cells at 48 and 72h vs control.

Finally, we analyzed differential gene expression in infected versus uninfected AMs obtained from COVID-19 ETA samples. Multiple interferon-stimulated genes (ISGs) were increased in infected compared to non-infected AMs in ETA samples (Fig. 4e), consistent with their being exposed to viruses more profoundly than neighboring cells. To ask whether ISG expression was a prominent feature of early infection, we returned to the PCLS system and compared uninfected, IAV, and SARS-CoV-2 exposure. We indeed found that AMs exposed to SARS-CoV-2 had upregulated ISGs. However, this induction was typically ~10-fold less in SARS-CoV-2, compared to IAV infection (Fig. 4f).

AMs are the sentinel immune cells in lung alveoli and relied upon to efficiently engulf and neutralize pathogens^16^. Here, we show that these cells are targeted by two of the most significant SARS-CoV-2 variants, leading to productive infection, viral propagation, and yet blunted interferon responses compared to IAV. These results suggest a depot effect where protective defenses are highjacked to facilitate viral production. AMs are migratory in the alveolar spaces^16^ and SARS-CoV-2-infected AMs could potentially spread infection to uninvolved areas of the lung leading to catastrophic viral loading of the lung, including as discussed in the accompanying manuscript, the loading of interstitial macrophage populations. We also found multiple macrophage subpopulations with SARS-CoV-2 viral reads, and it is certainly possible that these non-AM cells are also capable of productive infections. As ACE2 is an interferon-stimulated gene^17^, initial infection may be self-propagating, since our results indicated a critical dependency on ACE2 expression, although other viral entry mechanisms are potentially involved. As one recent example, in later stages of COVID-19 infection, and when antibodies have been generated, Fc gamma receptors appear to lead to monocyte infection^12^. In our model, the lack of antibodies likely rules out a role of Fc receptors in viral entry into AMs. It is intriguing to consider the long-lasting effects of this macrophage depot, which could be linked to non-resolving critical illness and long-term complications of COVID-19.

In summary, by examining the very early immune events in the human lung after SARS-CoV-2 infection, we discovered a specific tropism for lung myeloid populations and evidence of productive infection by AMs that has broad implications in unraveling the pathogenesis of severe SARS-CoV-2 infections in humans.

## The UCSF COMET Consortium

Cai Cathy, Bushra Samad, Suzanna Chak, Rajani Ghale, Jeremy Giberson, Ana Gonzalez, Alejandra Jauregui, Deanna Lee, Viet Nguyen, Kimberly Yee, Yumiko Abe-Jones, Logan Pierce, Priya Prasad, Pratik Sinha, Alexander Beagle, Tasha Lea, Armond Esmalii, Austin Sigman, Gabriel M. Ortiz, Kattie Raffel, Chayse Jones, Kathleen Liu, Walter Eckalbar, Billy Huang, Norman Jones, Jeffrey Milush, Ashley Byrne, Saherai Caldera, Catherine DeVoe, Paula Hayakawa Serpa, Eran Mick, Mayra Phelps, Alexandra Tsitsiklis, K. Mark Ansel, Stephanie Christenson, Gabriela K. Fragiadakis, Andrew Willmore, Sidney A. Carrillo, Alyssa Ward, Kirsten N. Kangelaris, Simon J. Cleary, Zoe M. Lyon, Vincent Chan, Nayvin Chew, Alexis Combes, Tristan Coureau, Kamir Hiam, Kenneth Hu, Billy Huang, Nitasha Kumar, Divya Kushnoor, David Lee, Yale Liu, Salman Mahboob, Priscila Munoz-Sandoval, Randy Parada, Gabriella Reeder, Alan Shen, Yang Sun, Sara Sunshine, Jessica Tsui, Juliane Winkler, Peter Yan, Michelle Yu, Shoshana Zha, Didi Zhu.

## Methods

### Precision Cut Lung Slices (PCLS)

Human donor lungs were obtained from Donor Network West. Lung lobes were inflated using 2% low-melting point agarose and incubated at 4°C^5^,^6^. After agarose consolidation, 1 cm^3^ lung tissue was placed on the precision compresstome VF-200 (Precisionary Instruments Inc.) for slicing. 300 μm slices were obtained and cultured in DMEM (UCSF Media Production), 1% penicillin/streptomycin (UCSF Media Production), and 10% fetal bovine serum (FBS) (Corning) in a 24-well plate.

### SARS-CoV-2 infections

Vero E6 and Vero-TMPRSS2 cells (gift from Dr. Melanie Ott) were cultured in DMEM supplemented with 10% FBS, penicillin/streptomycin, and L-glutamine (Corning) in a humidified incubator at 37°C and 5% CO2. SARS-CoV-2 virus (USA-WA1/2020 strain) was provided by Dr. Melanie Ott and propagated in Vero E6 or Vero-TMPRSS2 cells. SARS-CoV-2 B.1.617.2 (delta) variant was acquired from the California Department of Public Health, cultured in Vero-TMPRSS2 cells. For propagation, the Vero or Vero-TMPRSS2 cells were infected with the SARS-CoV-2 virus, incubated at 37°C, 5% CO2, and at 72h the supernatant was collected. The virus was aliquoted and stored at −80°C. All work was done under Biosafety Level 3 (BSL-3) conditions. Viral titer was quantified using a plaque assay in Vero cells^18^. Briefly, 10-fold dilutions of the virus stock were added to Vero cells in a 12-well plate for 1 hour, after which an overlay of 1.2% Avicel RC-581 in DMEM was added. The cells were incubated at 37°C, 5% CO2 for 96 hours. The cells were fixed with 10% formalin, stained with crystal violet, and washed with water. The plaques were counted to determine the titer of the virus stock.

### Influenza A virus (IAV) infections

As a virus of reference, we used influenza A/Puerto Rico/8/34 (*PR8*, H1N1) virus labelled with Venus to infect lung slices and BAL cells. PR8-Venus IAV was a gift from Yoshihiro Kawaoka (University of Wisconsin-Madison). The virus was produced in pathogen-free fertilized chicken eggs (Charles River) as published^19^. In brief, eggs were kept in an egg turner for 10 days. PR8-Venus IAV was injected into the allantoic cavity. Infected chicken embryos were incubated with the virus for 48 hours. Allantoic fluid was harvested, filtered, and snap-frozen in liquid nitrogen. Titers were determined with a hemagglutination assay. The work was done under BSL-2 conditions.

### Immunofluorescence imaging

After infection, PCLS were fixed in 4% PFA for at least 30 min. After saturation in PBS with 1% BSA, slices were stained with anti-CD45 (HI30, 304056, BioLegend), anti-spike (40150-R007, Sino Biological), anti-ACE2 (bs-1004R, BIOSS) anti-dsRNA (J2, 10010, Scicons) or anti-EpCAM-AF488 (9C4, 324210, BioLegend), for 1 hour in PBS with 1% BSA media at room temperature. PCLS were washed in PBS, counterstained with DAPI and attached to plastic coverslips using Vetbond (3M). Confocal imaging was performed using a Nikon A1R laser scanning confocal microscope with NIS-Elements software and a 16X LWD water dipping objective. Images were taken at more than 25 μm deep inside the slices to avoid the cutting artifact. 50 - 100 μm-thick images with a z-step of 1.5 μm were taken and analyzed using Imaris (Bitplane).

### Flow cytometry

At selected time points, PCLS were dissociated using DNase (4μg/ml) and Collagenase IV (200U/ml) at 37°C for 30min. Cells were filtered and stain for viability and surface markers (Supplementary Table 3). After fixation and permeabilization (BD cytofix/Cytoperm), cells were stained for spike and dsRNA (Supplementary Table 3). Data were collected using the BD LSRII Cytometer and analyzed using FlowJo version 10 (BD Biosciences).

### Bronchoalveolar lavage

A bronchoalveolar lavage (BAL) was done in a human lung lobe using ice-cold PBS. BAL cells were filtered and red blood cells lysed. BAL cells were plated in 24 well plates at 5 x 10^5^ cells per well in DMEM, 1% PS, 10% FBS containing 50 ng/ml of rhM-CSF (300-25, Peprotech). For selected experiments, BAL cells were treated for 2h before infection with an ACE2 blocking antibody at 10 μg/ml (AF933, R&D systems). SARS-CoV-2 was added to the cells at MOI 0.1 or 1. After 48h of infection, cells were recovered and stained for viability and surface markers (Supplementary Table 4). After fixation and permeabilization, cells were stained for intracellular spike expression. In selected experiments, AMs (live, EpCAM-, CD3-, CD19-, CD45+, HLA-DR+, CD169+) were flow-sorted prior to infection with SARS-CoV-2 using FACSAria Fusion (BD Biosciences).

### Virus replication assay

BAL cells were infected for 48h either with SARS-CoV-2 or IAV-Venus at MOI 0.1 or 1. At the end of the incubation, cell-free supernatant was recovered. This solution was used as inoculum for Vero E6 cells (SARS-CoV-2) or MDCK cells (IAV-Venus, gift from Michael A. Matthay). After 24 hours of incubation, cells were recovered and infection was assessed by flow cytometry.

### Single-cell RNA-sequencing

PCLS were dissociated as described above and dead cells were removed using Miltenyi Dead Cell Removal Kit (130-090-101, Miltenyi Biotec). For the multiplexing purpose, cells were then labeled with lipid modified oligonucleotides (LMO) and barcode oligos using the Multi-seq technique^20^. Cells were counted and the targeted cell number for loading was 8000 cells per sample. 10X encapsulation and library construction were done using Chromium Next GEM Single Cell 3’ Reagent Kits v3.1 per the manufacturer’s instruction. Multi-seq library preparation was done as previously described^20^. Libraries were mixed at an approximate 10:1 molar ratio of gene expression to LMO barcodes for sequencing. The sequencing was done on the Illumina NovaSeq 6000 using 10X Genomics recommended sequencing parameters.

### Data pre-processing of 10x Genomics Chromium scRNA-seq data

Sequencer-generated bcl data (Gene expression and Lipid Hashtag) was demultiplexed into individual fastq libraries using the mkfastq command on the Cellranger 3.0.2 suite of tools (https://support.10xgenomics.com). Feature-barcode matrices for all samples were generated using the Cellranger count command. Briefly, raw gene-expression fastqs were aligned to the GRCh38 reference genome annotated with Ensembl v85, and Lipid Hashtag fastqs were processed to count the incidences of each expected index per cell. Feature-barcode matrices were read into Seurat 4.0.1^21^ and poorly-captured genes (in < 3 cells) were dropped from the analyses. Matrices were further filtered to remove events with greater than 20% mitochondrial content, events with greater than 50% ribosomal content, or events with fewer than 100 total genes.

### Data quality control and normalization

The gene expression count matrices were normalized, and variance stabilized using negative binomial regression using the scTransform algorithm^22^ in the Seurat package. Cellular mitochondrial content, ribosomal content, and cell cycle state were regressed out of the data at this stage to prevent any confounding signal. The normalized matrices were transformed into a lower subspace using Principal Component Analysis (PCA) and 50 PCs per samples were used to generated Uniform Manifold Approximation and Projection (UMAP) visualizations. Cell clustering via expression was conducted using the Louvain algorithm. Cluster identities were assigned using Gene scores generated via the Seurat AddModuleScore function on a list of gene sets obtained from the Human Lung Cell Atlas.

### Demultiplexing of pooled single-cell libraries

In Lipid hashtagged libraries, the raw lipid tag counts were normalized using the Centered Log Ratio method (CLR) where HTO counts are divided by the geometric mean for that HTO across all cells and then log normalized. The resulting matrix was demultiplexes into donor samples using the Seurat HTODemux function^22^ using default parameters.

### Data integration and batch correction

scTransformed count matrices from all samples were integrated together using the in-built integration method provided by Seurat. The following commands were run in order, to integrate datasets ‘SelectIntegrationFeatures’ to identify shared “anchor” features for integration, ‘PrepSCTIntegration’ to subset objects based on identified anchor features, ‘FindIntegrationAnchors’ to identify anchor points between datasets based on the anchor features, and finally ‘IntegrateData’ to actually integrate the datasets based on the computed anchors. Downstream processing (PCA, UMAP, clustering and cluster assignment) was conducted similarly to the individual libraries.

### Endotracheal aspirates samples (ETA)

Endotracheal aspirate (ETA) samples were prospectively collected from seven adults requiring mechanical ventilation for the acute respiratory distress syndrome (ARDS) from COVID-19 as part of the COVID Multiphenotyping for Effective Therapies (COMET) study, as previously described^23^. Human participants were enrolled at two tertiary care hospitals in San Francisco, CA (UCSF Medical Center, Zuckerberg San Francisco General Hospital) under research protocol 20-30497 approved by the University of California San Francisco Institutional Review Board. ETA samples were collected within 1-40 days post-intubation for scRNA-sequencing. See Supplementary Table 2 for details.

### Statistical analyses

Statistical analysis was performed using GraphPad Prism v7.0e. For PCLS experiments (Fig. 1), significance was assessed using ANOVA with Sidak’s multiple comparison test. For Fig. 3 and Extended Data Fig. 5 (BAL cells and AM experiments), each dot represents the mean of 2 to 3 replicates from a single donor. Donors are color-coded across experiments. Paired multiple group comparisons (Fig. 3d and Extended Data Fig. 5g) were analyzed by one-way ANOVA with Dunnett’s multiple comparisons. Ratio paired t-test was used to compare two paired groups (ACE2 blockade, virus propagation assay, plaque assay).

## Acknowledgements

This work was supported by NIH funding as follows: NIAID-sponsored Immunophenotyping Assessment in a COVID-19 Cohort (IMPACC) Network [NIH/NIAID U19 AI1077439], R35 HL140026 (C.S.C.), 3P01AI091580-09S1 (JPR), R01 AI052116 (M.F.K.), R01 AI160167 (M.R.L.), R35 HL161241 (M.R.L.), P30 DK063720 (UCSF Flow Cytometry CoLab), and Shared Instrument Grant 1S10OD021822-01 (UCSF Flow Cytometry CoLab). We also acknowledge funding from COVID-19 Fast Grants (M.O. and M.R.L.), UCSF Clinical and Translational Science Institute (CTSI), and the Bakar UCSF ImmunoX Initiative. M.O. thanks the Rodenberry foundation, Pam and Ed Taft, the Innovative Genomics Institute, and the Gladstone Institutes for their support.

## Author contributions

All authors contributed to manuscript preparation. Conceptualization: MM, RY, MFK, MRL. Experimentation: MM, RY, LR, CRS, LH, JPR. Human samples: The UCSF COMET Consortium, CL, PGW, DJE, CMH, CSC, MAM, CRL. Data analysis: MM, RY, AAR, RTD, KHH, MFK, MRL. Reagents: AS, MO. Supervision: MFK, MRL. Manuscript writing: MM, RY, AAR, MFK, MRL.

---

**Supplementary Table 1.**
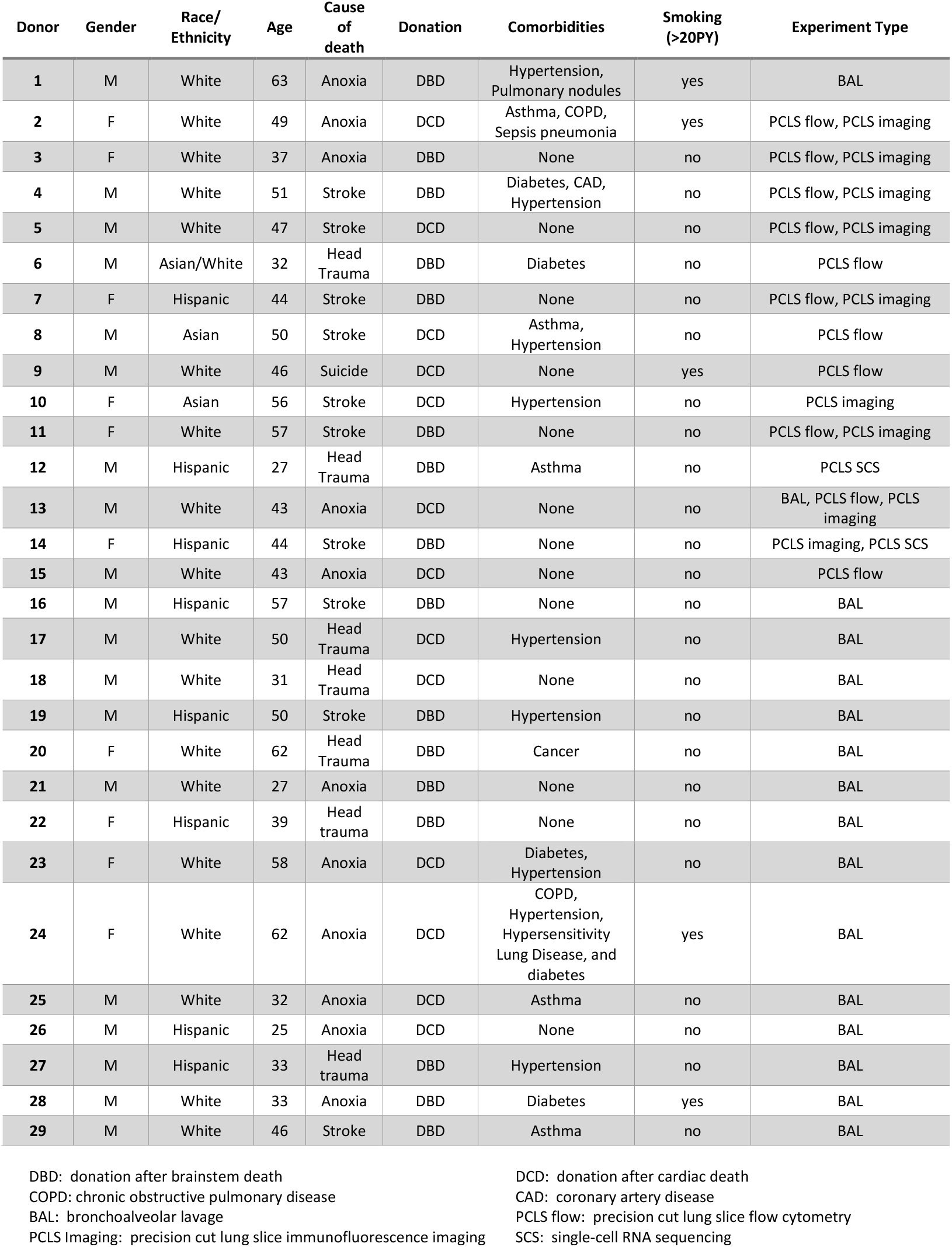
Human lung donor demographics and experimental assignments.

**Supplementary Table 2.** Demographic and clinical information for COVID-19 ETA samples.

| Subject | Age | Gender | Ethnicity | ARDS (Y/N)<br>Berlin<br>definition | ETA<br>sampling<br>after<br>intubation<br>(days) | P/F ratio at<br>time of ETT<br>sampling | ICU LOS<br>(days) | Hospital<br>LOS (days) | Death (Y/N) |
| --- | --- | --- | --- | --- | --- | --- | --- | --- | --- |
| 1 (1) | 34 | F | White | Yes | 0 | 180 | 12 | 14 | No |
| 2 (365) | 55 | M | White | Yes | 40 | 110 | 35 | 60 | No |
| 3 (414) | 58 | M | Other /<br>Multiple<br>Races | Yes | 1, 3 | 165 | 19 | 27 | No |
| 4 (415) | 62 | F | Black /<br>African<br>American | Yes | 2 | 124 | 14 | 21 | No |
| 5 (419) | 68 | M | Other /<br>Multiple<br>Races | Yes | 7 | 175 | 26 | 50 | No |
| 6 (476) | 59 | M | White | Yes | 12 | 160 | 44 | 78 | No |
| 7 (389) | 66 | F | White | Yes | 2 | 144 | 39 | 55 | No |

ARDS: acute respiratory distress syndrome
ETA: endotracheal tube aspirate
P/F ratio: $\text{PaO}_2/\text{FiO}_2$
ICU LOS: intensive care unit length of stay
Hospital LOS: hospital length of stay

**Supplementary Table 3.** Flow cytometry panel for PCLS experiments.

| **Staining** | **Antibody (clone)** | **Lot number** | **Dye** | **Catalog number (Supplier)** |
| --- | --- | --- | --- | --- |
| **Surface**<br>**Staining** | CD169 | 1621398 | AF594 | FAB5197T (R&D Systems) |
|  | ACE2 | BJ07067522 | PE | bs-1004R (BIOSS) |
|  | EpCAM | B250368 | BV650 | 324226 (BioLegend) |
|  | CD31 (WM59) | 8232937 | BV605 | 562855 (BD Biosciences) |
|  | CD45 (HI30) | B333796 | BV421 | 304032 (BioLegend) |
|  | CD14 (M5E2) | B275828 | BV711 | 301838 (BioLegend) |
|  | CD3 (SK7) | 0314461 | BB700 | 566575 (BD Biosciences) |
|  | CD19 (SJ25C1) | 1069967 | BB700 | 566396 (BD Biosciences) |
|  | Zombie (viability) | B331984 | NIR | 77184 (BioLegend) |
|  | HLA-DR (G46-6) | 1266420 | BUV395 | 564040 (BD Biosciences) |
| **Intracellular**<br>**Staining** | Spike | 1619059 | AF647 | FAB105805R (R&D Systems) |
|  | dsRNA (J2) | J2-2007 | AF488 | 10010 (Scicons) |

**Supplementary Table 4.** Flow cytometry panel for BAL experiments.

| **Staining** | **Antibody** | **Lot number** | **Dye** | **Reference** |
| --- | --- | --- | --- | --- |
| **Surface**<br>**Staining** | CD169 | 1621398 | AF594 | FAB5197T (R&D Systems) |
|  | ACE2 | BJ07067522 | PE | bs-1004R (BIOSS) |
|  | CD15 (W6D3) | 1089776 | BV786 | 741013 (BD Biosciences) |
|  | CD16 (3G8) | B321940 | BV605 | 302040 (BioLegend) |
|  | CD45 (HI30) | B333796 | BV421 | 304032 (BioLegend) |
|  | CD14 (M5E2) | B275828 | BV711 | 301838 (BioLegend) |
|  | CD3 (SK7) | 0314461 | BB700 | 566575 (BD Biosciences) |
|  | CD19 (SJ25C1) | 1069967 | BB700 | 566396 (BD Biosciences) |
|  | Zombie (viability) | B331984 | NIR | 77184 (BioLegend) |
|  | HLA-DR (G46-6) | 1266420 | BUV395 | 564040 (BD Biosciences) |
| **Intracellular**<br>**Staining** | Spike | 1619059 | AF647 | FAB105805R (R&D Systems) |
|  | IFITM3 (EPR5242) | GR3416716-1 | AF488 | Ab198559 (Abcam) |

