## Extended Data Figures for "Immediate myeloid depot for SARS-CoV-2 in the human lung"

Extended Data Figure 1

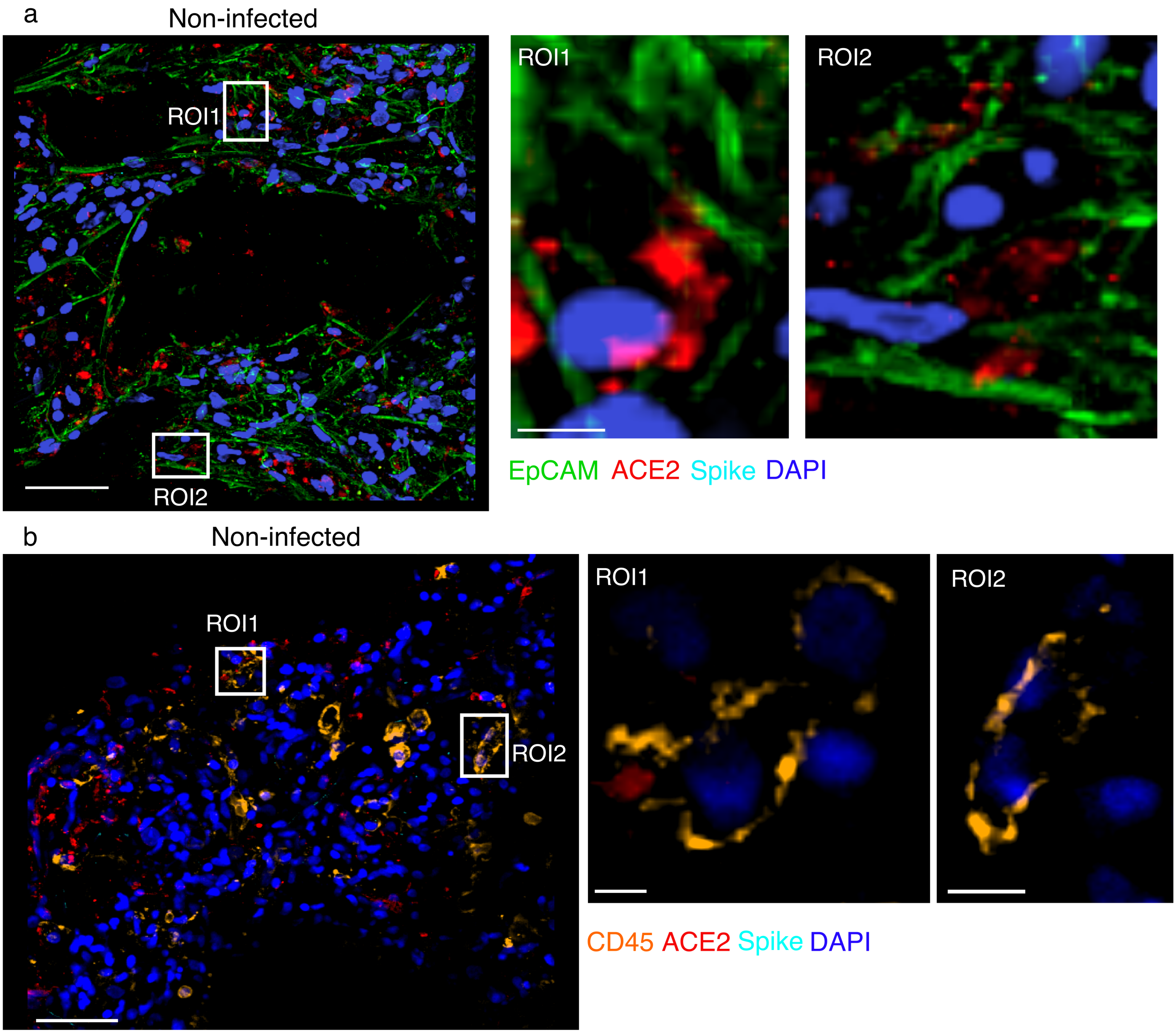


**Extended Data Figure 1. ACE2 expression in epithelial and immune cells in non-infected PCLS.** Confocal images were obtained from non-infected controls (scale bar = 50 µm). Zoom area is marked by the white rectangles. For each zoomed area, slices appear on the side (scale bar = 5 µm in (**a**) and 10 µm in (**b**). (**a**) PCLS stained for EpCAM (green), ACE2 (red) and spike (light blue). (**b**) PCLS stained for CD45 (orange), ACE2 (red) and spike (light blue).

Extended Data Figure 2

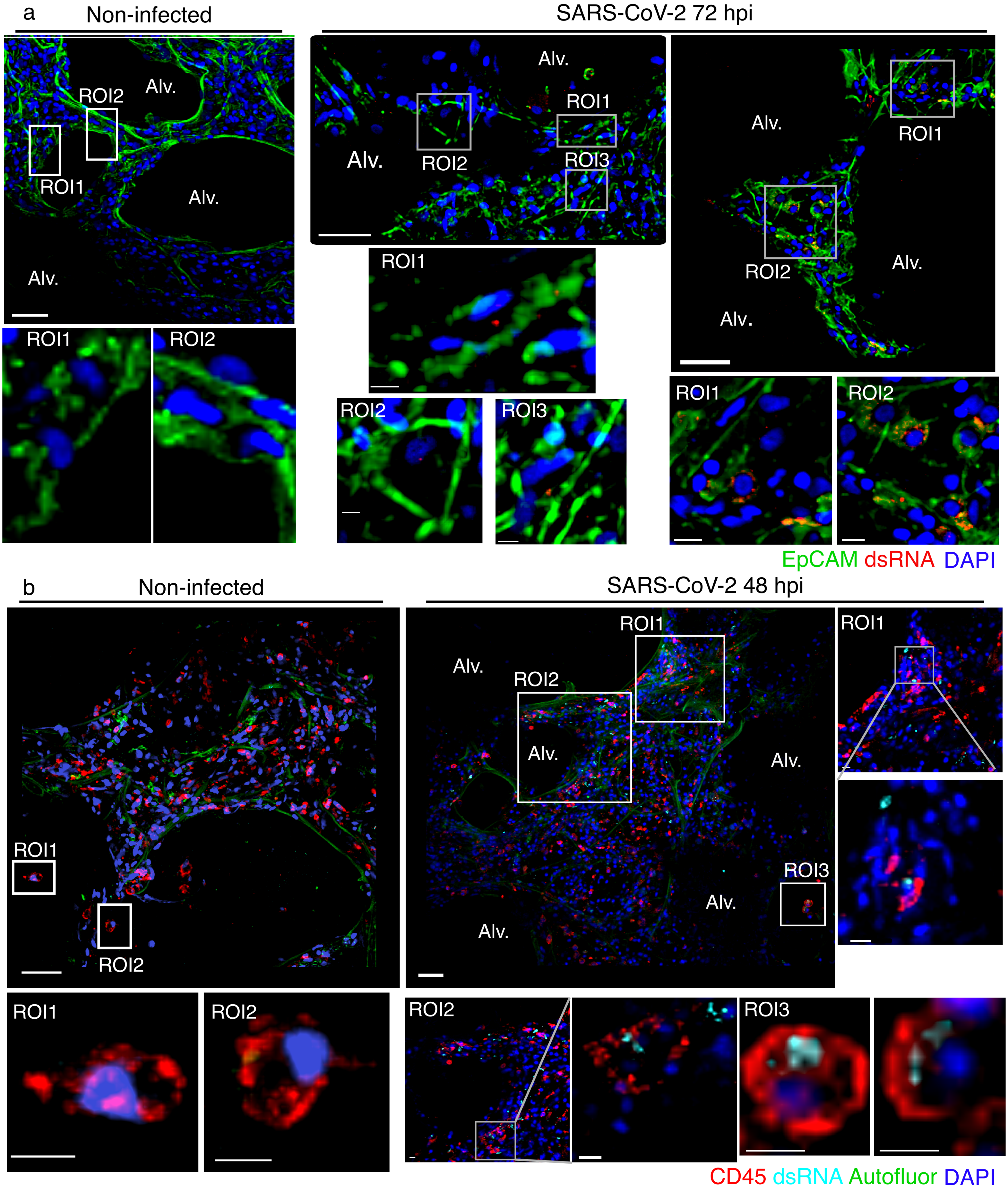


**Extended Data Figure 2. SARS-CoV-2 infection drives dsRNA production in both epithelial and immune cells in PCLS.** PCLS were infected with SARS-CoV-2 and used for confocal imaging. Alveolar spaces (Alv.) are indicated in the large image (scale bar = 50 µm). Zoom area is marked by the white rectangle. For each zoomed area, z-stacks appear on the side (scale bar = 10 µm). (**a**) PCLS stained for EpCAM (green) and dsRNA (red). (**b**) PCLS stained for CD45 (red) and dsRNA (light blue). Autofluorescence appears in green.

### Extended Data Figure 3

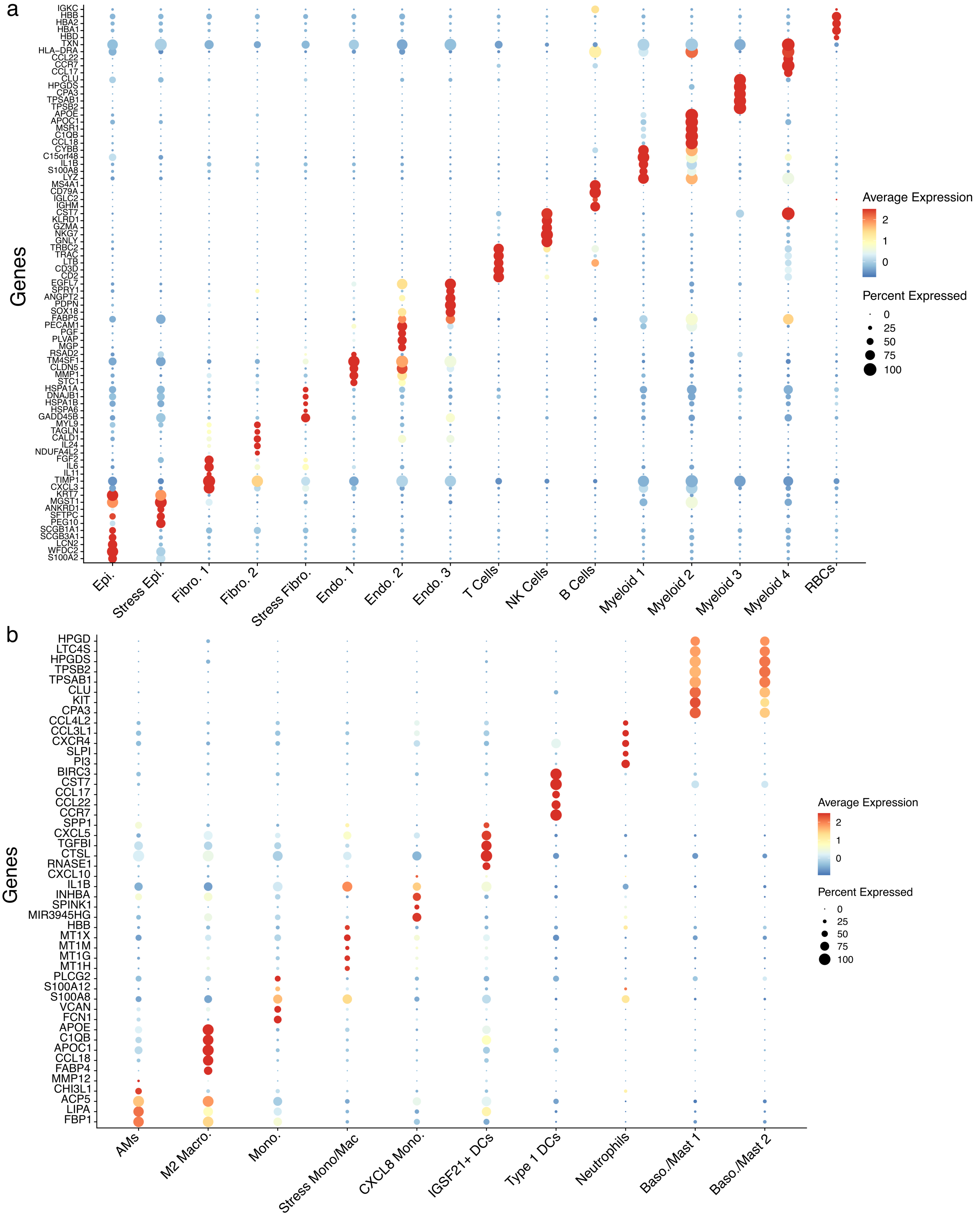


**Extended Data Figure 3. Cell type annotations for scRNA-seq in human PCLSs.** (**a**) A dot plot describing the expression of the top 5 genes per identified cluster in the human PCLS dataset, used to assign cell type annotations. (**b**) A dot plot describing the expression of the top 5 genes per identified myeloid subcluster in the human PCLS dataset, used for myeloid cell type annotations.

Extended Data Figure 4

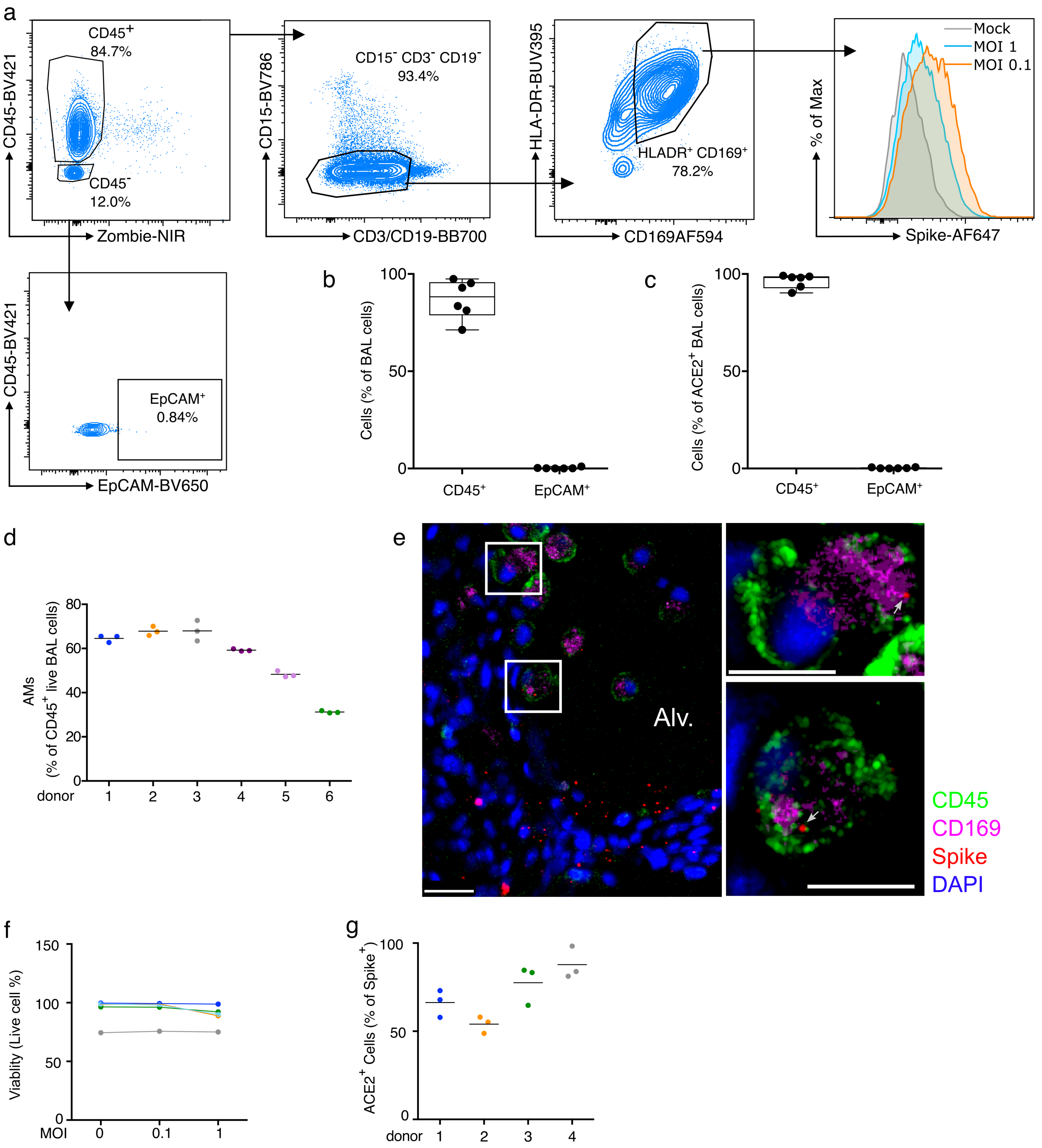


**Extended Data Figure 4. Human BAL cell profile and effects of SARS-CoV-2 infection.** (**a**) Flow cytometry gating of human BAL cells. (**b**, **c**) Proportions of leukocytes (CD45+) and epithelial cells (CD45- EpCAM+) in total and ACE2+ BAL cells (n=6). (**d**) Proportion of AMs (CD45+ CD15-, CD3- CD19- HLA-DR+ CD169+) amongst total, live BAL cells (n=6). (**e**). PCLSs were infected with SARS-CoV-2 and stained for CD45 (green), CD169 (violet) and spike (red) before confocal imaging. Alveolar space (Alv.) is indicated in the large image (scale bar = 30 μm). Zoom areas are marked by the white rectangle (scale bar = 15 μm). (**f**) Viability of BAL cells infected for 48h with SARS-CoV-2 at MOI 0.1 or 1 or control (n=5). (**g**) ACE2 expression in spike+ AMs (n=4).

Extended Data Figure 5

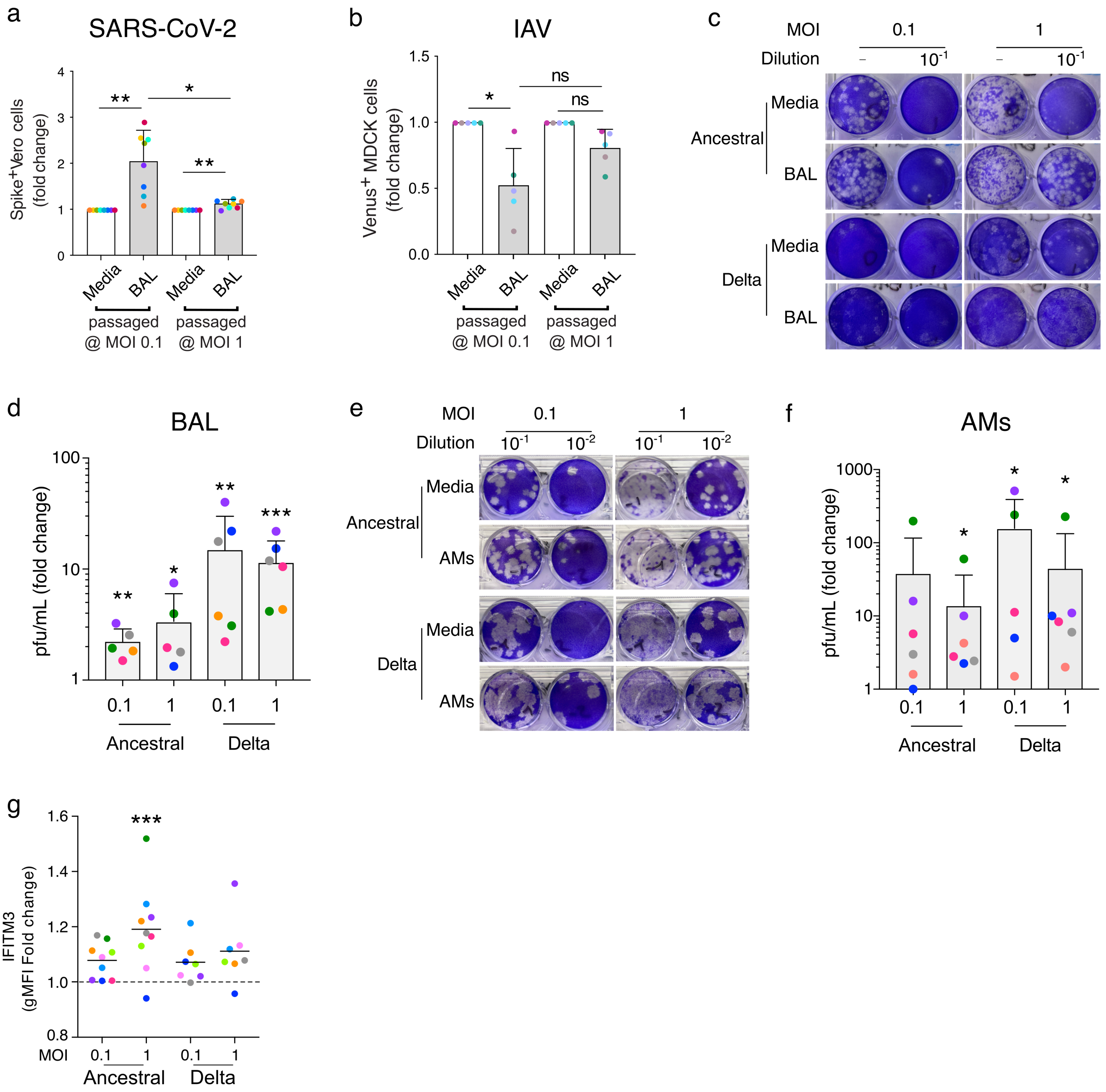


**Extended Data Figure 5. Infection of BAL cells and flow-sorted AMs with SARS-CoV-2 variants.** (**a**) BAL cells were infected with SARS-CoV-2 for 48h at MOI 0.1 and 1. As a control, SARS-CoV-2 was incubated in cell culture media alone for 48h. Cell-free supernatant was used to infect VeroE6 cells. At 24h after Vero E6 infection, cells were stained for intracellular spike expression. Using flow cytometry, infected Vero E6 percentages was determined (n=8, mean ± SD). Results appear in fold change compare to media incubation. (**b**) Similarly, IAV-Venus was used to infect BAL cells (MOI 0.1 and 1) or incubated with media for 48h. Cell-free supernatant was used to infect MDCK cells. At 24h after MDCK infection, cells were recovered for flow cytometry (n=5, mean ± SD). Venus expressing cells percentage was measured and appears as a fold change from media incubation. (**c**, **d**) Viral titer was determined by plaque assay following 48h incubation of BAL cells and SARS-CoV-2 viruses. (**d**) Results appear as fold change from media incubated virus (n=5-6, mean ± SD). (**e**, **f**) BAL AMs (live, EpCAM-CD45+CD3-CD19-HLA-DR+CD169+) were flow-sorted and SARS-CoV-2 variants were incubated either with AMs (MOI 0.1 - 1) or media for 48h. Cell free supernatant was used for plaque assay. (**f**) Results are expressed as fold-change compared to media incubation (n=5-6, mean ± SD). (**g**) IFITM3 staining in AMs after infection with SARS-CoV-2 (Ancestral, delta) at MOI 0.1 - 1 (n=7-9). Results appear in fold change compare to uninfected cells. *p<0.05, **p<0.01, ***p<0.001.

### Extended Data Figure 6

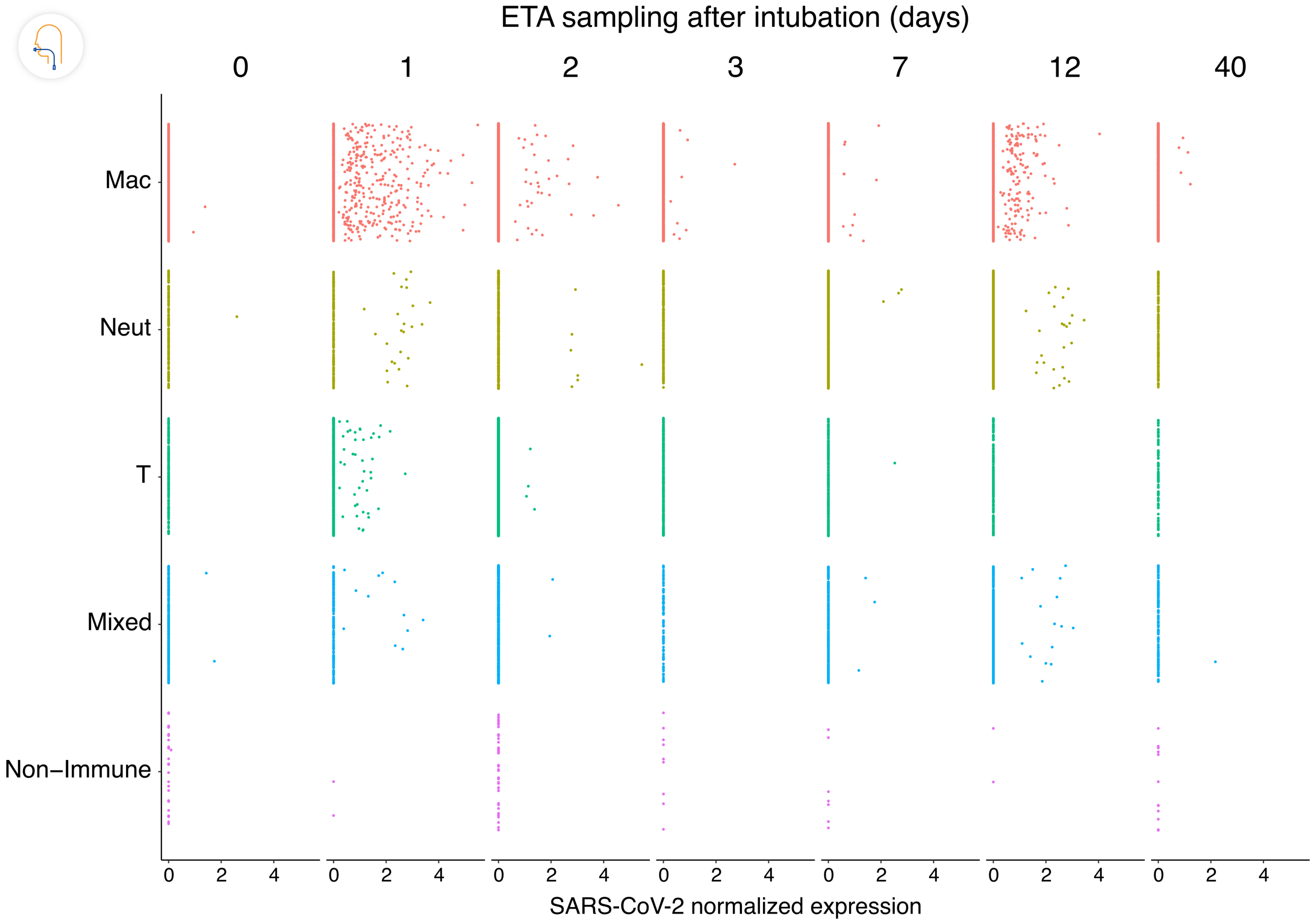


**Extended Data Figure 6. Distribution of per-cell normalized SARS-CoV-2 expression in landmark cell types in human ETA samples from COVID-19 ARDS subjects.** The days (D0-D40) at the top of the graph represent the days post-intubation when the ETA samples were collected. Each column represents a unique COVID-19 subject except for post-intubation day 2, where 2 subjects are represented. See Supplementary Table 2 for details.

### Extended Data Figure 7

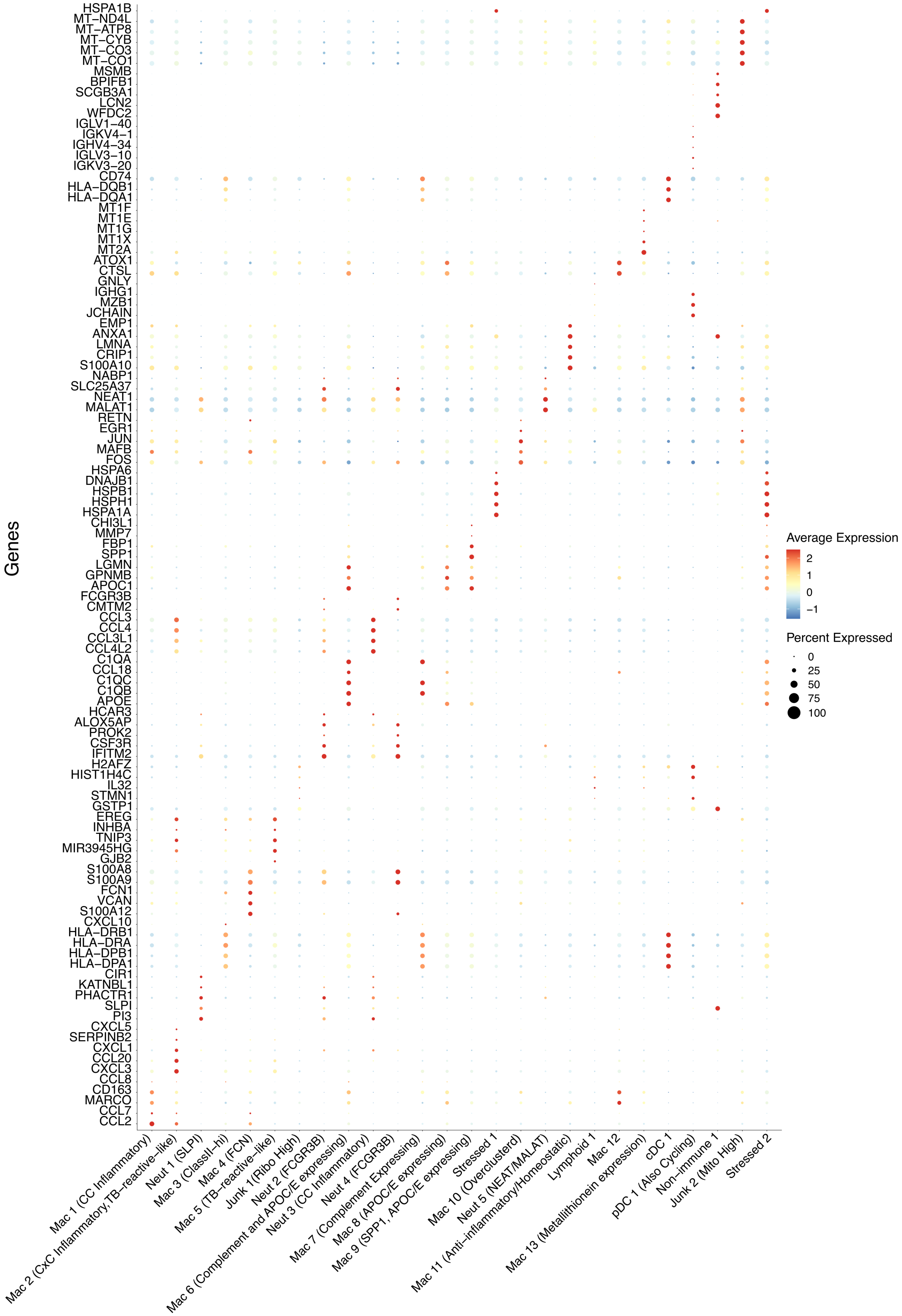


**Extended Data Figure 7. Cell type annotations for scRNA-seq in human ETA samples.** A dot plot describing the expression of the top 5 genes per identified cluster in the human ETA dataset, used to assign cell type annotations.
